# Expectation Shapes Neural Preparation for AI-generated and Real Image Processing: Evidence from EEG

**DOI:** 10.64898/2026.07.17.738592

**Authors:** Yonghao Chen, Anna Eiserbeck, Martin Maier, Felix Klotzsche, Simon M. Hofmann, Julia Baum, Birgit Nierula, Anna Hilsmann, Sebastian Bosse, Arno Villringer, Rasha Abdel Rahman, Michael Gaebler, Vadim V. Nikulin

## Abstract

AI-generated images, videos, and news have become an inseparable part of daily life, leading to increasing skepticism toward online information. Understanding how expectations about the presence of AI-generated content influence perception is crucial for elucidating the underlying neural mechanisms and facilitating the generation of naturalistic avatars. In this study, we analyzed an existing EEG dataset (N = 29) in which participants viewed emotional facial photographs of only real individuals while being informed that upcoming faces were either real (“REAL”) or AI-generated (“FAKE”). We examined pre-stimulus oscillatory activities in the one-second EEG interval before stimulus onset and found significantly lower alpha power when participants expected “FAKE” compared to “REAL” faces. This effect was region-specific, particularly in right occipito-parietal and temporo-parietal regions, as identified by both sensor- and source-level analyses. In addition, the modulation effect between pre-stimulus alpha activity and post-stimulus event-related potentials (ERPs) was measured. A significant correlation between changes in late positive potential (LPP) amplitudes and pre-stimulus alpha power was observed exclusively for “FAKE” smiling faces, consistent with our previous findings. These results suggest that AI-related expectations modulate neural preparatory states, with lower alpha activity presumably reflecting increased attentional demands for stimuli believed to be artificially generated. This study demonstrates that top-down beliefs systematically shape both pre- and post-stimulus neural dynamics, providing new insights into how cognitive expectations bias perceptual processing, with implications for understanding human-AI interaction and improving the design and evaluation of AI-generated content in real-world contexts.

## Introduction

Generative models have great potential to improve our daily life and work efficiency in various domains. One impactful capability is generating naturalistic avatars and videos with minimal human effort (Pataranutaporn et al., 2021; Wahab, 2025; Wu et al., 2025). Specifically, deepfakes and other media generated by generative models are now widespread on major social platforms (Menczer et al., 2023; Boediman, 2025; Birrer & Just, 2025). However, users are increasingly exposed to large amounts of misinformation, including “fake news” and AI-generated images and videos (Beridze et al., 2019; Westerlund, 2019; Fan et al., 2025; Folorunsho et al., 2025), which leads to growing skepticism toward content on social media even if it is actually made by humans (Whittaker et al., 2020; Altay & Gilardi, 2024; Clark et al., 2026; Dunn et al., 2026). Accumulating evidence suggests that humans find it hard to visually distinguish between AI-generated photos and real human photographs (Nightingale & Farid, 2022; Miller et al., 2023; Dunn et al., 2026). Despite growing research interest in the perception of AI-generated faces within the field of neuroscience, most existing studies have focused primarily on the perception of fake images or videos themselves (Moshel et al., 2022; Tarchi et al., 2023; Chen et al., 2024; Becker et al., 2026); relatively few studies systematically explore the neural processing underlying our top-down beliefs about whether an image is real or AI-generated (Eiserbeck et al., 2023). Understanding the cognitive and neural mechanisms underlying this top-down modulation effect is critical not only for improving the quality of AI-generated content but also for a broader scientific understanding of the societal impact of the AI revolution.

Predictive coding is a widely accepted theory stating that the human brain is an adaptive system, continuously updating its internal models based on prior experience to minimize prediction errors during cognition (Rao & Ballard, 1999; Friston & Kiebel, 2009; Huang & Rao, 2011). Accordingly, whether perceived content is labelled as AI-generated would fundamentally bias their perceptual processing of an image. Empirical evidence supports this notion across different domains. Studies suggest that repeated interactions with robots reduce the overall sense of “eeriness” caused by the uncanny valley effect (Złotowski et al., 2015), and individual differences in the tendency to attribute mental states to robots can be decoded from users’ resting-state brain activities (Bossi et al., 2020). Similarly, individuals more frequently exposed to digital characters evaluate computer graphics (CG) faces more positively than people who report less frequent exposure (Di Natale et al., 2024). All those findings suggest that experience with and personal attitude toward virtual or synthetic avatars would indeed bias perceptual and cognitive processes. Moreover, another study found that labeling videos as AI-generated reduced both their perceived accuracy and participants’ willingness to share them, even if the content is real or human-made (Altay & Gilardi, 2024). The gap between these behavioral phenomena and the underlying neural processes remains unresolved.

Neural oscillations, particularly oscillations in the alpha frequency band (8-13 Hz), are frequently associated with top-down processes under the framework of predictive coding (Arnal & Giraud, 2012; Samaha et al., 2015; Alamia & VanRullen, 2019; Turner et al., 2023). Specifically, alpha activity typically increases when the brain suppresses irrelevant sensory input (Klimesch et al., 2007; Jensen & Mazaheri, 2010; Compton et al., 2019).

Within this context, increased alpha power serves as a neurophysiological marker of cortical inhibition, effectively reducing neural excitability in regions processing irrelevant inputs to prioritize goal-directed predictions. Conversely, alpha desynchronization is typically associated with increased excitability (Pfurtscheller et al., 1996; Başar et al., 1997; Klimesch et al., 2007; Jensen & Mazaheri, 2010; Clayton et al., 2018). Despite the established role of alpha oscillations in predictive and attentional processes, it remains unclear whether oscillatory activity can also serve as a reliable biomarker of neural responses to presumed AI-generated content.

In recent work (Eiserbeck et al., 2023), we used electroencephalography (EEG) to track neural signals from 30 participants who were asked to evaluate photographs of human faces displaying three emotions (happy, neutral, angry), with each image preceded by a cue labeling it as either real or AI-generated (deepfakes). To avoid confounds from bottom-up visual differences and isolate the effects of prior expectation, all stimuli were in fact photographs of real people. The EEG analysis focused on event-related potentials (ERPs) and showed that presumed real smiling faces elicited canonical emotion effects with differences relative to neutral faces on three ERP components (P100, N170, Early Posterior Negativity; EPN). In contrast, smiling faces presumed to be AI-generated showed none of these early effects. Furthermore, only the ’fake’ smiling faces elicited enhanced Late Positive Potential (LPP) activity, suggesting a shift toward more effortful evaluation. Notably, negative facial expressions induced typical neural emotion effects regardless of whether they were labeled as “REAL” or “FAKE” before stimulus presentation. While these findings demonstrate that top-down expectations modulate post-stimulus neural responses, the role of pre-stimulus oscillatory activity remains unclear. Specifically, oscillatory dynamics occurring between the presentation of instruction cues (“REAL” or “FAKE”) and stimulus onset may play an important role in shaping subsequent neural responses. Therefore, the present study aims to examine how pre-stimulus oscillations depend on expectation toward the upcoming stimuli and how these oscillations modulate subsequent ERPs.

Our study was preregistered (https://osf.io/4x6k8). We hypothesized that pre-stimulus oscillatory activity reflects the manipulated belief whether an image is a real photograph or an AI-generated face, resulting in a difference in power spectrum and spatial distributions of oscillations across one or some specific frequency bands (Theta, Alpha and Beta). Furthermore, we expected that these baseline activities would exert a top-down influence on post-stimulus processing, specifically modulating emotion-related ERP components like the LPP or EPN. Likewise, we anticipated that this modulation effect would be particularly evident in the processing of smiling faces, in agreement with the earlier research. Overall, this study aims to provide novel insights into how top-down expectations about whether the upcoming stimulus is AI-generated or not shape neural preparation for face perception.

## Methods

### Participants

The analysis utilized data of 29 participants (21 female, 8 male, *M*_age_ = 25.97, *SD*_age_ = 5.04) out of the original 30 participants (Eiserbeck et al., 2023), as one participant was excluded due to excessive muscle artifacts in the EEG recordings during the pre-stimulus interval. All participants had normal or corrected-to-normal vision and provided written informed consent before their inclusion in the study. The study was conducted according to the principles expressed in the Declaration of Helsinki. It was approved by the ethics committee of the Department of Psychology at Humboldt-Universität zu Berlin. Participants received either course credit or monetary compensation of €10 per hour.

### Stimuli

Real photographs of 180 unique individuals (90 female, 90 male) were selected as stimulus materials from three face databases: Radboud Faces Database (Langner et al., 2010), Karolinska Directed Emotional Faces (Lundqvist et al., 1998), and FACES (Ebner et al., 2010). To control facial features, the face stimuli were cropped into an oval shape, excluding hair, ears, and neck. Although participants were informed that some images were AI-generated (experimental manipulation of belief), all stimuli were in fact real photographs. More details are presented in Eiserbeck et al.(2023).

### Experimental Procedure

After signing informed consent, participants received a verbal introduction (based on a standardized text) on the increasingly high quality of AI-generated faces and watched a silent video that illustrated the creation of “fake” faces using Generative Adversarial Networks (GANs). The introduction and video highlighted the potential of these models and noted that AI-generated faces are often indistinguishable from real ones (for full video and verbal script details, see Eiserbeck et al., 2023, Supplementary Information). Participants were then informed that they would see both real and AI-generated facial images during the formal experiment, with each picture preceded by the corresponding label “REAL” or “FAKE”.

The experiment consisted of two parts: a free viewing task and a facial expression rating task, which were always carried out in this exact order. These tasks differed only in their response demands: participants were required to provide a rating after each stimulus in the second task, but not in the first.

The rating task was divided into two blocks, where each of the 180 face stimuli was presented once per block in a random order. Each face was preceded by a “REAL” or “FAKE” cue, resulting in a total of 360 trials (60 trials per individual Emotion × Information condition). The assignment of “REAL” and “FAKE” information to each face was counterbalanced across participants. Short breaks were provided every 30 trials to mitigate fatigue. **Figure 1** shows the exact procedure of the rating task. Trials began with a 500-ms fixation cross, followed by the cue “REAL” or “FAKE” (500 ms). A blank screen was then presented for a random duration of 500-1000 ms prior to face onset. While the exact pre-stimulus duration for each trial was not logged, the total interval between the word cue and stimulus onset ranged from 1000 to 1500 ms. Subsequently, the respective face stimulus was displayed for 800 ms. Following a 500-ms post-stimulus delay (1300 ms after face onset), a continuous 101-point rating scale (values 1–101) appeared for facial expression judgments. The scale endpoints were labeled “very negative” and “very positive,” with the scale direction being counterbalanced across participants. Participants utilized a mouse-controlled slider to indicate their valence judgment. The scale remained on-screen until a response was recorded. Each trial concluded with a 1000 ms inter-trial interval (ITI) before the start of the next sequence. The free viewing task, which preceded the rating task, contained the same number of trials and followed the same structure, except that no rating was required.

**Figure 1:**
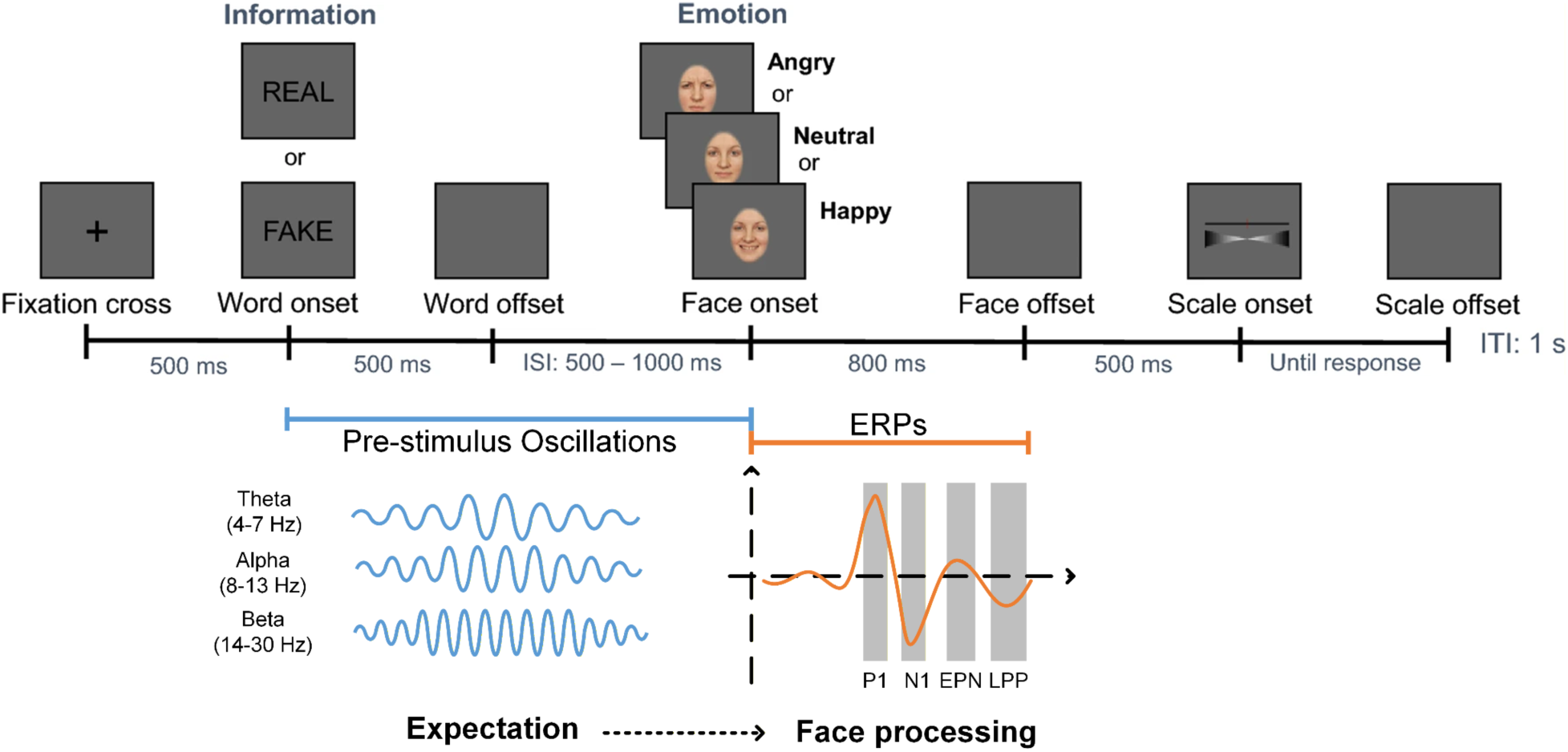
Schematic representation of the trial sequence and the primary EEG features extracted. Each trial began with a fixation cross followed by a word cue (“REAL” or “FAKE”) and the subsequent presentation of an emotional face (angry, neutral, or happy). Participants then provided a valence rating for each expression using a 101-point scale ranging from “very negative” to “very positive.” ISI = Inter-stimulus interval; ITI = Inter-trial interval. The example face image was taken from Eiserbeck et al., 2023.

After the experiment, participants filled in a questionnaire about demographic information and the experiment. Finally, a formal debriefing was conducted to clarify that all facial stimuli stemmed from real photographs and that the “REAL/FAKE” designations were part of the experimental manipulation.

### EEG Recording and Preprocessing

The EEG was recorded using a Brain Products BrainCap with Ag/AgCl electrodes at 64 scalp sites (63 EEG + 1 EOG) according to the extended 10–20 system, at a sampling rate of 500 Hz, with all electrodes referenced to the left mastoid. During recording, low- and high-cutoff filters (0.016 Hz and 1000 Hz) were applied, and electrode impedances were maintained below 10 kΩ. A full description of the recording setup is provided in Eiserbeck et al. (2023). EEG data were preprocessed and analyzed using MATLAB R2025a (The MathWorks Inc., Natick, USA) and the EEGLAB toolbox (version v2024.2, Delorme & Makeig, 2004). Given that the main purpose of this study is different from the previous study, the EEG preprocessing pipeline was adjusted accordingly as follows:

- Applying a high-pass filter at 1Hz to remove DC offset, baseline drifts, and slow-wave artefacts.
- Removing bad electrodes based on manual visual inspection of each participant’s raw data and on the automated channel-rejection algorithms (*_pop_rejchan*) provided by the EEGLAB toolbox. The removed bad channels were interpolated using a spherical spline method (Perrin et al., 1989).
- Applying independent component analysis to remove noise. The resulting independent components were automatically classified using the ICLabel plugin within the EEGLAB toolbox (Pion-Tonachini et al., 2019). Components identified as eye artefacts, muscle activity, or channel noise with a classification probability higher than 0.9 were labeled as noise and excluded from the data.
- Finally, all EEG electrodes were re-referenced to a common average reference.

### Oscillations Analysis (Spectral Power)

Pre-stimulus oscillatory activity was analyzed within a fixed time window of -1000 ms to 0 ms relative to face stimulus onset. This fixed-window approach was implemented because the precise inter-stimulus interval (ISI) could not be calculated due to the absence of word-onset triggers. Consequently, our analysis did not focus on time-locked phase information with respect to “FAKE” vs “REAL” stimuli; instead, we utilized the power spectral density (PSD) as the primary metric to quantify the intensity of pre-stimulus oscillations. For each trial, the PSD value was calculated as the normalized squared magnitude of the fast Fourier transform (FFT) output after applying a Hamming window. Using a pre-stimulus interval that closely precedes the face presentation has the advantage of capturing the neural state immediately before the stimulus onset. Spectral power was analyzed in three primary frequency bands of interest: Theta (4-7 Hz), Alpha (8-13 Hz), and Beta (14-30 Hz). Mean PSD values were calculated by averaging the power within these frequency ranges across the entire 1000 ms pre-stimulus interval. The difference in PSD between the two conditions was examined using non-parametric cluster-based permutation tests (Maris & Oostenveld, 2007), as well as a typical right occipitotemporal channel (PO8), which has been proven crucial in face-relevant studies (Pitcher et al., 2011; Rossion, 2014). Data from both tasks were concatenated separately for each condition, yielding 360 trials per condition; single-trial PSD values were calculated and subsequently averaged.

### ERP Analysis

To ensure consistency with the analytical framework established in Eiserbeck et al. (2023), we maintained a similar pipeline for ERP analysis to replicate the results, except for the pre-processing approaches mentioned above. However, unlike Eiserbeck et al. (2023), the present study included data from both tasks (free-viewing & rating task) in all analyses to increase statistical power for oscillation analysis, and one participant was removed in the screening stage. Consequently, these methodological adjustments may introduce minor discrepancies in our findings compared to the original study. A low-pass filter at 40Hz was applied to remove high-frequency noise that did not contribute significantly to evoked responses in face perception (Zhang et al., 2024). Data were segmented into epochs of [-200 ms,1000 ms] relative to face stimulus onset and baseline-corrected using the 200 ms pre-stimulus interval. The amplitudes of four ERP components (P100, N170, EPN, LPP) were separately calculated based on different pre-selected channels and time windows used in Eiserbeck et al.(2023):

**P100 (P1)**: P100, an early component peaking around 100 ms after stimulus onset, was analyzed as a marker of the initial stages of face perception (Eimer, 2000; Rossion, 2014). Five typical parieto-occipital electrode sites (O1, O2, Oz, PO7, PO8) were selected, and the amplitudes were acquired by averaging the activities within the time window of [80ms, 130ms] relative to stimulus onset.
**N170**: N170, a typical ERP component associated with face-specific processing, was analyzed as a marker of the early structural encoding phase in face perception (Bentin & Deouell, 2000; Rossion & Jacques, 2011). Eight parieto-occipital electrode sites (TP9, TP10, P7, P8, PO9, PO10, O1, O2) were selected, and the time window for averaging was [130 ms, 200 ms] relative to stimulus onset.
**EPN**: EPN (early posterior negativity) was analyzed as a marker of the early emotional processing of faces (Schupp et al., 2004; Schindler et al., 2021). Six posterior electrode sites (PO7, PO8, PO9, PO10, TP9, TP10) were selected, and the time window for averaging was [200 ms, 350 ms].
**LPP**: LPP (late positive potential), a sustained ERP component peaking during the later stages of stimulus processing (Blechert et al., 2012; Bublatzky et al., 2014), was analyzed as an index of sustained attention and evaluative processing, reflecting the perception of realism and deeper cognitive appraisal of emotional facial expressions in this study. Six centro-parietal electrode sites (Pz, Cz, C1, C2, CP1, CP2) were selected, and the time window for averaging was [400 ms, 600 ms].

Emotion effects (*A_instr_*_,*emo*_) were evaluated by computing the difference in ERP amplitudes between emotional facial stimuli (Happy & Angry) and neutral stimuli:

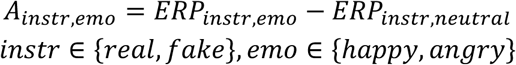

Where the *ERP_instr_*_,*emo*_ represents the averaged ERP amplitudes for a specific condition and emotional expression of stimulus, four ERP components (P100, N170, EPN, LPP) were calculated separately. *emo* represents the emotional expressions of facial stimuli (“happy” or “angry”), and *instr* represents the conditions of instructions (“REAL” or “FAKE”). The differences between the two conditions were calculated by:

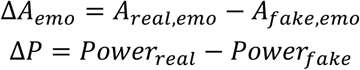

Where *Power* represents the mean power of pre-stimulus oscillatory activities for both conditions (“REAL” or “FAKE”) within the same electrode cluster utilized for calculating ERP amplitudes, the Spearman correlation coefficients between Δ*A_emo_* and Δ*P* were calculated to evaluate how the expectation of AI-generated content modulates the processing of emotional faces.

### Source Localization

To identify the specific brain regions underlying the observed effects, Linearly Constrained Minimum Variance (LCMV) beamforming (Van Veen et al., 1997) was applied to reconstruct power differences within the source space. The regularization parameter was set at 5% for inversion of the covariance matrix to ensure numerical stability. The forward model was acquired by using the Colin 27 head model (Holmes et al., 1998) and the boundary element method (BEM, Mosher et al., 1999). Inverse modelling was computed separately per participant and condition before it was averaged for each condition across all subjects. To make sure that the reconstructed source power was directly comparable between “REAL” and “FAKE”, a common beamform filter was calculated for each participant by concatenating trials from all conditions. Eight electrodes (IO1, A2, F9, F10, TP9, TP10, PO9, PO10) were excluded from source reconstruction for all participants due to strong muscle artefacts or because they failed to align precisely with the geometry of the head model. The inverse models based on the power spectrum were calculated separately for individuals, and the results were averaged across all participants within each condition. Anatomical classification was based on Brodmann areas (Brodmann, 2006). All source-space analyses were implemented in MATLAB R2025a using the FieldTrip Toolbox (Oostenveld et al., 2011).

### Behavioral Data Analysis

In addition to the EEG data, we analyzed a behavioral variable that was recorded after the experiment: the deliberate inclusion score, i.e., participants’ own evaluation of the extent to which they deliberately or consciously considered the “REAL”/ “FAKE” information when judging the facial expressions. The score was rated using a continuous scale ranging from 1 (“Not at all”) to 101 (“Very strongly”). Specifically, the correlation between the deliberate inclusion score and the magnitude of power difference of pre-stimulus oscillatory activities between the two conditions was investigated.

### Statistical Analysis

To compare oscillatory power and ERP amplitudes across different experimental conditions, paired *t*-tests were implemented. To correct the statistics for multiple comparisons (e.g., comparing alpha power in multiple electrodes between “REAL” versus “FAKE” simultaneously), we used the non-parametric cluster-based permutation tests (Maris & Oostenveld, 2007), i.e., randomly shuffling the condition labels 1000 times to construct a null distribution of cluster-level statistics. Only clusters in the original data that exceeded the 95% percentile of the permutation distribution (corresponding to *p-*value < 0.05) were considered statistically significant.

The Spearman correlation was chosen as the main analytical approach in all the correlation analyses mentioned in this study to ensure robustness against the influence of outliers (De Winter et al., 2016). For all analyses, the statistical significance level was set to *p*-value < 0.05. All statistical analyses were performed in MATLAB R2025a.

## Results

### Spectral power

Spectral analysis revealed a classic *1/f*-shaped spectrum in the pre-stimulus interval with a pronounced peak in the alpha frequency range (**Figure 2**). The subsequent analysis revealed that despite the pre-stimulus neural activities exhibiting comparable spatial patterns across the two conditions, with maxima over occipito-parietal areas, the magnitude of the power differed between the “REAL” and “FAKE” conditions within the alpha frequency band (8-13 Hz). In contrast, no significant differences were observed in other frequency bands (*p* > 0.05), suggesting that the observed preparatory effect was specific to alpha oscillations. Significant electrode clusters were detected by applying cluster-based permutation tests (*p* < 0.01, 1000 permutations). Although alpha activity was most prominent in occipital regions as expected, the clusters showing significant differences between the two conditions spanned two distinct topographical regions: a right occipito-parietal cluster and a left temporal cluster.

**Figure 2:**
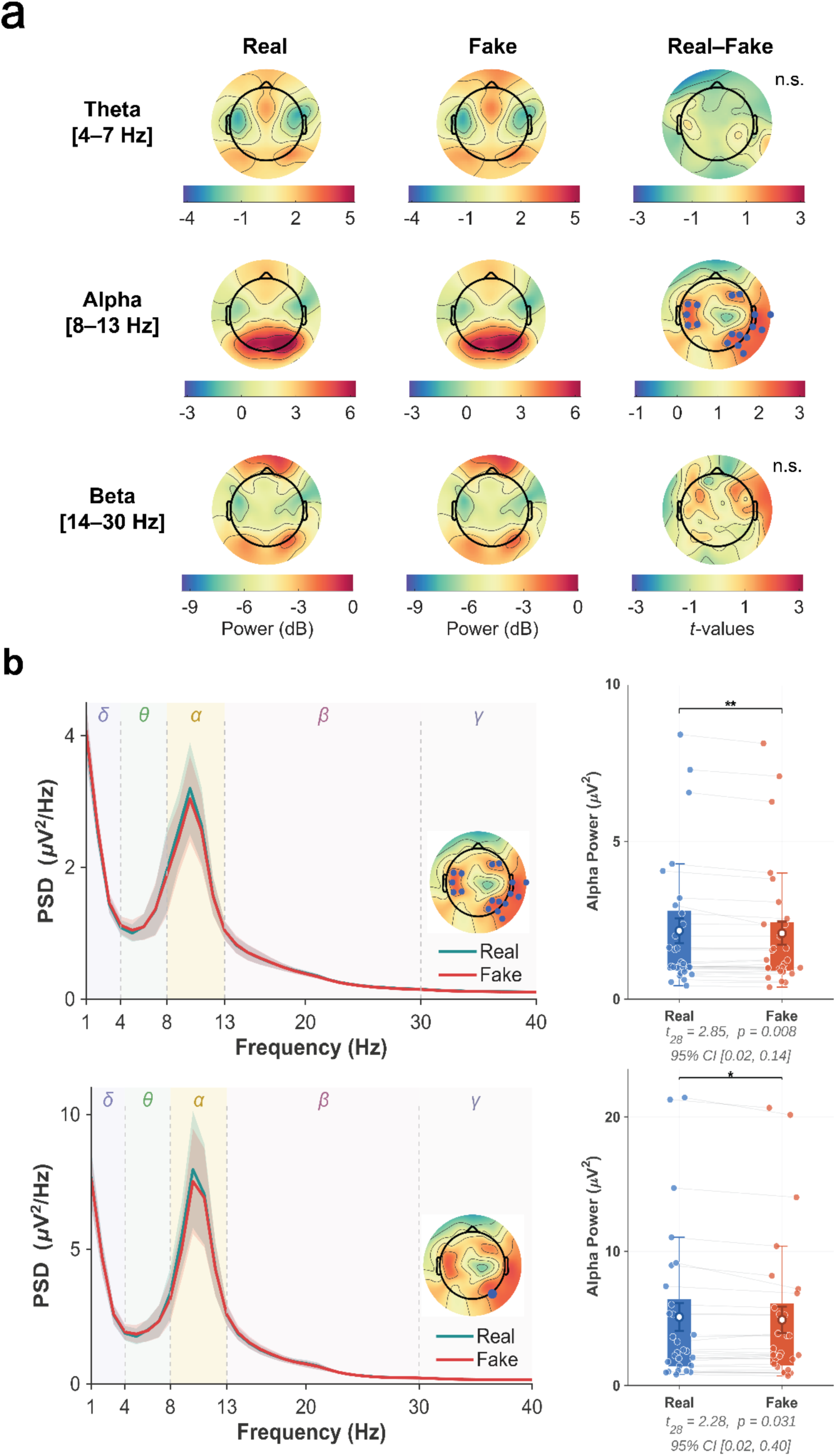
Topographies and differences in pre-stimulus oscillatory activities. (a) Averaged topographical distribution of power across three frequency bands (Theta, Alpha, Beta) and the power comparisons between the “REAL” and “FAKE” conditions (*t*-values of paired *t*-test). Significant clusters are indicated by blue dots. (b) PSD plots and the statistical results of alpha-band power differences for the identified clusters and the typical electrode PO8 (*: *p* < 0.05, **: *p* < 0.01). The colored dots represent individual participant alpha power for the respective conditions; the white dots represent the group means. Error bars indicate 95% confidence intervals.

In general, weaker alpha activity was present when the participants expected to see AI-generated faces, evidenced by both clusters (*p* < 0.01, *t* (28) = 2.85) and one typical electrode, PO8 (*p* = 0.031, *t* (28) = 2.28). Notably, while the power of the oscillations varied by condition, the alpha peak frequency remained stable for both conditions. For both group-level averages and individual participants, the peak frequency was centered at 10 Hz.

### Source analysis

Source reconstruction results further underlined the sensor space analysis (**Figure 3**). Reconstructed sources for both “REAL” and “FAKE” were primarily localized to the occipital and parietal regions. Differences in power magnitude were most pronounced in angular gyrus (BA 39) and superior temporal gyrus (BA 22) in the right temporo-parietal junction (rTPJ), as well as associative visual cortex (BA 19) in the left hemisphere.

**Figure 3:**
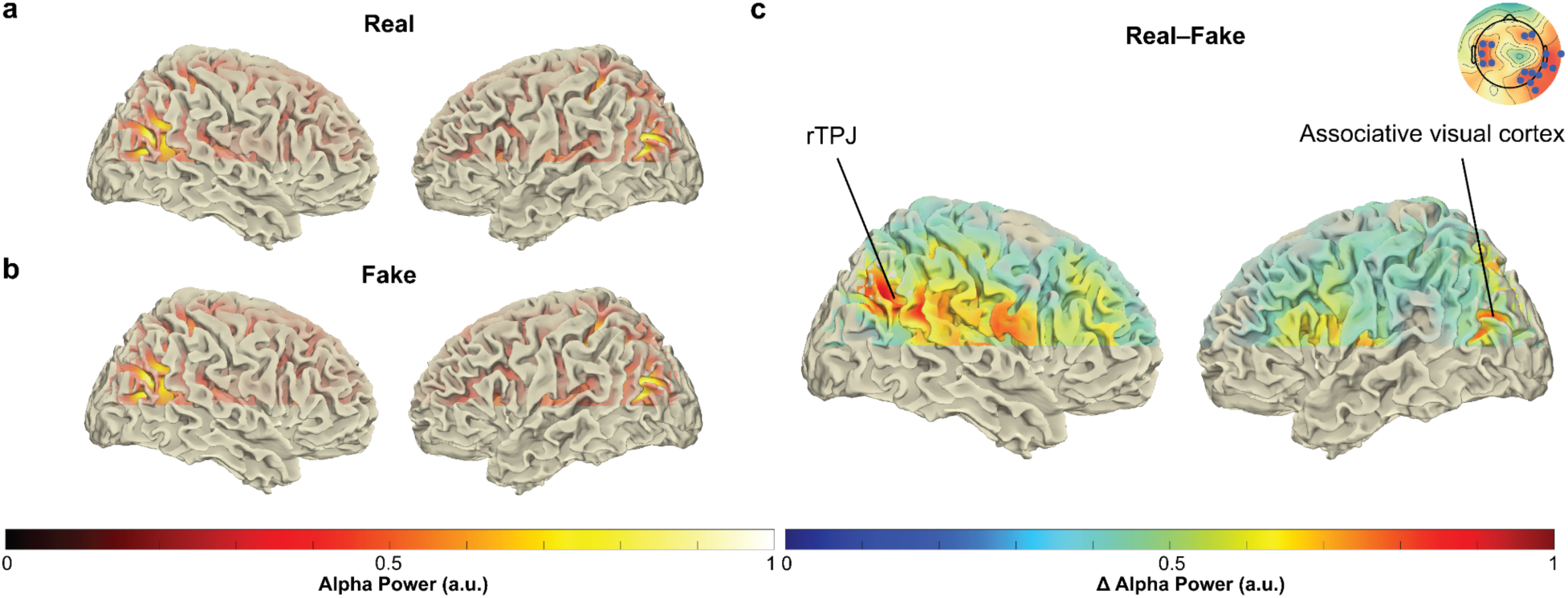
Source-localized pre-stimulus alpha activity. (a) Mean alpha power in the “REAL” condition. (b) Mean alpha power in the “FAKE” conditions. (c) Mean difference in alpha power between conditions (“REAL” minus “FAKE”). The topography plot shows the significant clusters in the sensor space analysis. All plots represent data after min-max normalization.

### ERP

ERP results were similar to Eiserbeck et al. (2023): presumed “FAKE” angry faces evoked similar neural activity to “REAL” faces (*A_fake_*_,*an*g*ry*_ ≈ *A_real_*_,*an*g*ry*_). In contrast, the neural differentiation between happy and neutral expressions of presumed “FAKE” faces was attenuated compared to “REAL” ones (*A_fake_*_,ℎ*appy*_ > *A_real_*_,ℎ*appy*_ ≈ 0) for two components (N170, and EPN). However, different from Eiserbeck et al. (2023), there are no significant amplitude differences that emerged during the other two processing stages (*A_fake_*_,ℎ*appy*_ ≈ *A_real_*_,ℎ*appy*_ > 0 for P1 & LPP), this might be due to the changes in the preprocessing stage and incorporating data from both tasks. More details regarding the ERP waveforms and statistical results are presented in the **Supplementary**.

The relationship between the difference in power of pre-stimulus alpha activity (Δ*P*) and the corresponding shifts in emotional ERP amplitudes (Δ*A*) is illustrated in **Figure 4**. Only the differential LPP amplitude for happy faces was significantly and positively correlated with pre-stimulus alpha oscillations (*ρ* = 0.49, *p* < 0.01); this indicates that participants with higher alpha power differences between the “REAL” and “FAKE” conditions also exhibited more pronounced LPP modulation in response to happy faces. Other, earlier, ERP components (P100, N170, and EPN) were not significantly linked to pre-stimulus alpha activity, but showed a trend toward a negative relationship.

**Figure 4:**
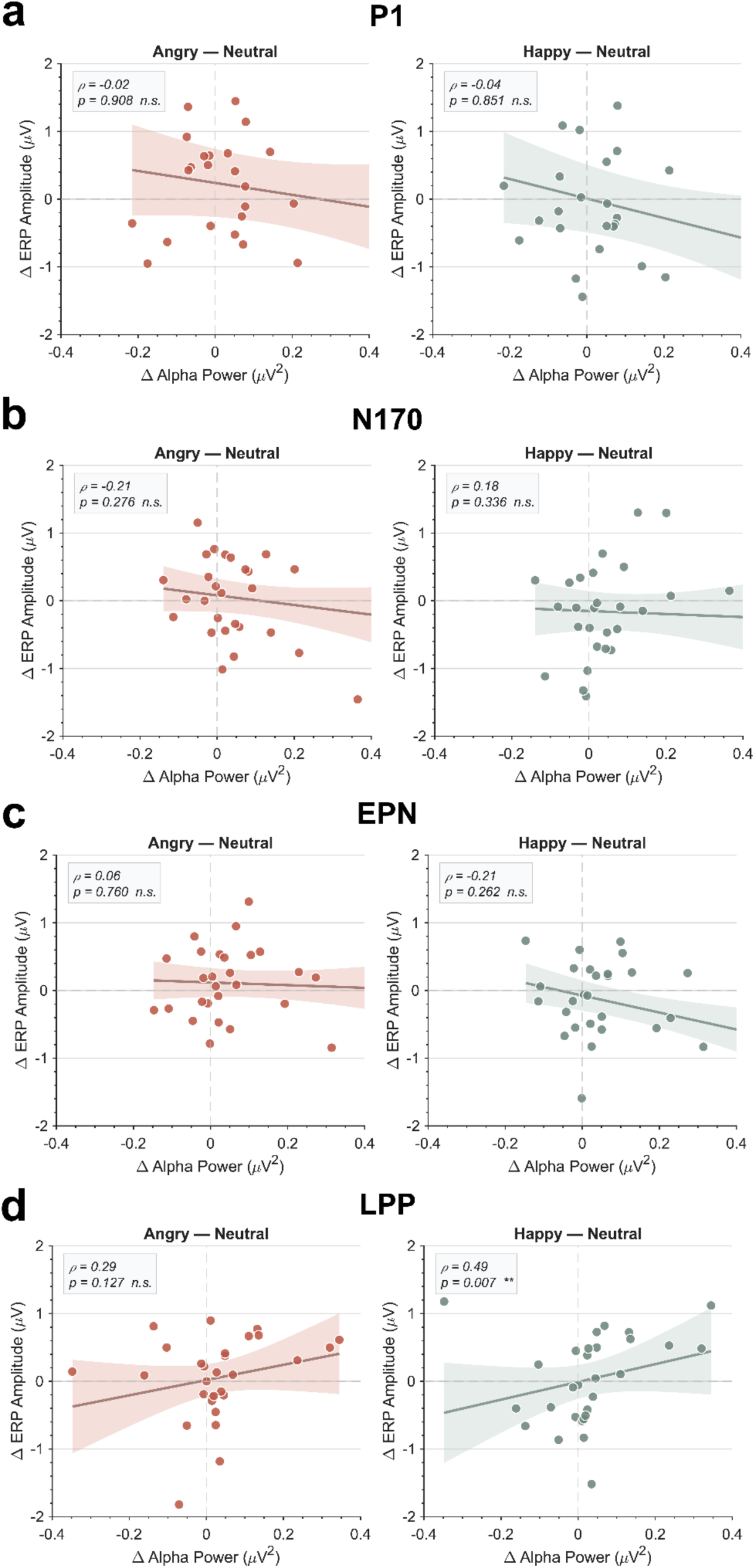
Correlation analysis between pre-stimulus alpha power modulation and ERP responses. The scatter plots illustrate the relationship between the differential alpha power and the emotional arousal effects for four distinct ERP components: (a) P1/P100, (b) N170, (c) EPN, (d) LPP. Differential ERP amplitudes (Δ*A*, Y-axis) and alpha power (Δ*P*, X-axis) were calculated by contrasting the “REAL” and “FAKE” instructional conditions as described in the Methods section. Solid lines indicate the linear regression fit; *ρ* denotes the Spearman correlation coefficient, with *p* representing the associated statistical significance (*: *p* < 0.05, **: *p* < 0.01, n.s.: *p* > 0.05).

### Behavioral results

As shown in **Figure 5**, the deliberate inclusion score showed a negative correlation with the pre-stimulus alpha power difference (*ρ* = -0.37, *p* = 0.049). This inverse correlation indicates that participants who reported a higher degree of conscious, deliberate reliance on the “REAL” vs. “FAKE” labels exhibited a smaller magnitude of difference in pre-stimulus alpha activity between the two conditions.

**Figure 5:**
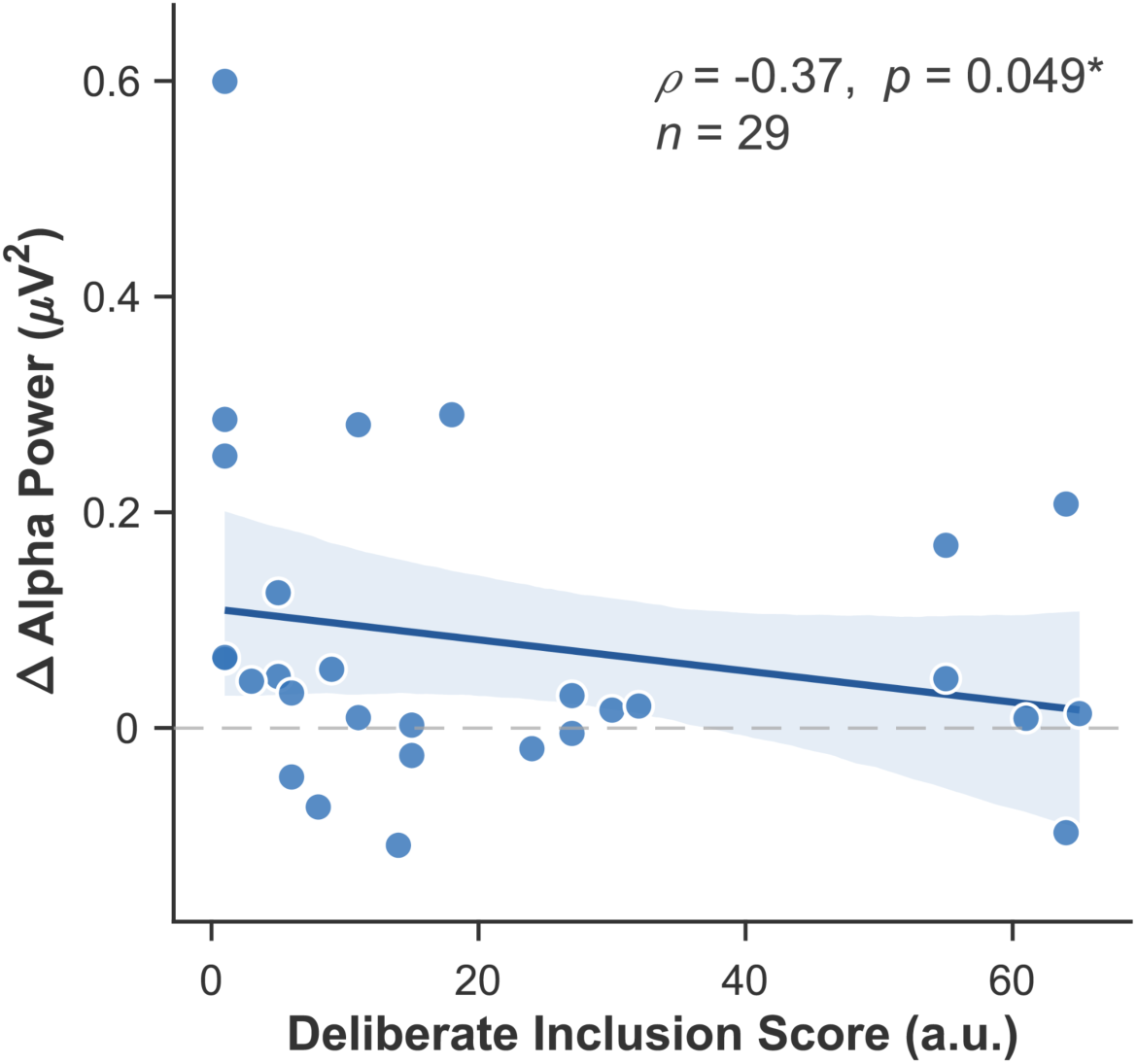
Correlation analysis of deliberate inclusion scores versus alpha power differences (“REAL” minus “FAKE”, *: *p* < 0.05). X-axis indicates the deliberate inclusion score for each participant; the Y-axis represents the mean alpha power extracted from the significant clusters described above.

## Discussion

This study investigated how expecting to see either real or AI-generated faces biases the perception and evaluation of real human faces at both behavioral and neural levels. We found that this effect can be tracked through changes in the brain oscillations. Specifically, we observed a significant attenuation of alpha activity when participants anticipated AI-generated faces. Meanwhile, the oscillations in other frequency bands (theta and beta) remained largely unchanged. Additionally, our analysis revealed that these pre-stimulus alpha shifts were correlated with the difference in post-stimulus LPP amplitudes, while the earlier ERP components (P1, N170, and EPN) showed no correlation. Together, these findings suggest that although expectations regarding AI-generated faces influence both early and late stages of face processing, as evidenced by ERP modulations, pre-stimulus alpha oscillations specifically predict the extent of expectation-related effects on later evaluative processing indexed by the LPP.

Why did this effect occur, and why only for alpha oscillations? Many previous studies have suggested that alpha attenuation serves as a marker of increased cortical excitability (Pfurtscheller et al., 1996; Başar et al., 1997; Klimesch et al., 2007; Jensen & Mazaheri, 2010; Clayton et al., 2018; Stephani et al., 2021; Stephani et al., 2025). Similarly, we propose that the expectation of encountering an AI-generated face serves as a task-relevant modulator of cortical readiness to process visual stimuli. Given that “REAL” human faces are the “default” social stimulus, the “FAKE” label likely transitioned participants from a passive viewing state to an active, more evaluative state, resulting in alpha attenuation. Another possible explanation is that the observed alpha desynchronization may reflect an affective bias caused by the “FAKE” label. Participants are likely to show unconscious skepticism or negative associations toward AI-generated content, even when merely informed through labeling (Altay & Gilardi, 2024). This state of distrust likely triggers increased emotional arousal, which is consistently associated with a decrease in alpha-band power (Hofmann et al., 2021; Schubring & Schupp, 2021; Codispoti et al., 2023). Specifically, this is consistent with the findings that only the LPP amplitude changes for happy faces were correlated with shifts in pre-stimulus alpha power. For angry faces, the label-induced arousal may have been congruent with the negative emotional content of the stimulus, producing a relatively strong response across conditions and thereby reducing the sensitivity to detect condition-specific modulation. In contrast, for happy faces, the mismatch between negative anticipatory bias and positive emotional content may have allowed individual differences in pre-stimulus alpha activity to exert a stronger influence on later evaluative processing, as reflected in the LPP.

This selective correlation pattern is particularly noteworthy because Eiserbeck et al. (2023) demonstrated that the same instructional manipulation influenced multiple stages of face processing. Specifically, they found that presumed deepfake smiles showed no canonical emotion effects in P1, N170, and EPN components, whereas presumed real smiles did. This suggests that while the belief about AI-generated content influences both early and late processing stages, pre-stimulus alpha power may specifically predict individual differences in late-stage evaluative processing rather than early perceptual encoding. This dissociation could indicate that pre-stimulus alpha reflects preparatory cognitive states related to evaluative rather than perceptual mechanisms, even though the experimental manipulation (“REAL”/“FAKE” labels) affects the entire temporal sequence of face processing. Furthermore, the present findings can be interpreted within a predictive coding framework, in which the brain continuously generates and updates predictions to minimize prediction error (Rao & Ballard, 1999; Huang & Rao, 2011). In this context, expectancy-related influences would emerge primarily at later processing stages, when a mismatch arises between the prior belief that a face may be AI-generated and the fact that the actual sensory input is a real human face, thereby selectively modulating late components such as the LPP. This implies that the detection of realism could be more closely associated with the late stage of face perception, and the participant might unconsciously judge the realism of faces even when their expectations have been experimentally manipulated.

The topographical characteristics of the pre-stimulus oscillations were examined using both sensor-level analyses and source localization in this study. Consistent with prior literature on resting-state dynamics, theta and alpha oscillations exhibited typical spatial patterns, with theta activity centered over fronto-midline regions (Ishihara & Yoshii, 1972; Klimesch, 1999; Karakaş, 2020) and alpha activity predominantly localized to parietal– occipital areas (Berger, 1929; Klimesch et al., 2007; Jensen & Mazaheri, 2010). In contrast, beta activity did not exhibit sensorimotor distribution over central areas at the resting state, which may reflect task-related preparatory activity. Notably, differences in pre-stimulus alpha power were primarily localized in two regions: an extended cluster spanning the angular gyrus and superior temporal gyrus within the right occipito-parietal and temporo-parietal regions, and the associative visual cortex in the left hemisphere. The cluster in the right hemisphere may reflect fluctuations in brain states related to social cognition, particularly within the right temporo-parietal junction (rTPJ), which has been widely implicated in social cognition and attribute belief to other people within the framework of theory-of-mind (Saxe et al., 2004; Aichhorn et al., 2009; Saxe & Kanwisher, 2013). These results suggest that interactions with AI-generated avatars may engage similar neural structures as those involved in typical social interactions with real humans, particularly within core social-cognitive networks. However, processing presumed AI-generated images may require greater attentional or cognitive effort, leading to differences in the magnitude of neural responses rather than substantial differences in the spatial distribution of activated regions. Secondly, the ROIs in the left associative visual cortex may, in addition, reflect interactions between visual perception and stored semantic knowledge, since the left associative visual cortex is closely linked to the ventral visual stream, which is involved in processing object identity and semantic information associated with visual stimuli (Schneider, 1969; Albright, 2012; Collins & Olson, 2014).

Accordingly, these results may reflect modulation driven by participants’ conceptual or semantic processing of the visual cues (“REAL” & “FAKE”).

Moreover, we also investigated the impact of individual-level differences by correlating the deliberate inclusion score and differences in pre-stimulus alpha power. The results suggested that participants who more deliberately considered realism labels exhibited reduced differences in alpha power between the two conditions. These findings support the idea that alpha desynchronization reflects changes in cortical excitability, as both conditions may elicit a more similarly activated brain state when participants allocate more attentional resources to processing the instructions. In contrast, when processing is more implicit, the FAKE label—representing a deviation from the default expectation of real human faces—triggers stronger alpha desynchronization, resulting in larger condition differences. However, the results only narrowly reached the conventional threshold for statistical significance (*p* = 0.049), which might be due to the limited sample size in this study (*N* = 29). Replication in larger samples, ideally combined with more detailed measures of participants’ beliefs and attitudes toward AI-generated content, will be helpful for assessing the reliability and generalizability of the present findings.

Our study systematically tested how experimentally manipulated expectations about viewing AI-generated faces influence late-stage neural processing. There are some limitations and open questions that could be addressed in future research. First, all stimuli applied in this study were photographs of real human faces; however, recent research suggests that the human brain may indeed be sensitive to differences when viewing state-of-the-art AI-generated images, as reflected in the corresponding neural activity patterns (Tarchi et al., 2023; Beckmann et al., 2024). A possible direction for future research would be to apply a passive viewing paradigm in which AI-generated and real human faces are presented, followed by participants’ subjective judgments of realism. Whether similar effects—reflected in differences in magnitude and spatial patterns shown in this study— can also be observed in event-related desynchronization (ERD) within such a paradigm remains to be further investigated. Secondly, to what extent prior personal experience with AI-generated content affects the results observed in this study remains unknown. In particular, given that the largest differences were observed within the rTPJ, it would be valuable to further examine whether individual factors such as age, sex, and social experience modulate this neural pathway. Last but not least, the present analysis mainly focused on cumulative pre-stimulus activity. Future studies could extend this work by applying time–frequency analyses in designs with a fixed interval between instruction and stimulus onset, which may provide further insight into the underlying neural mechanisms.

Taken together, this study offers new insights into the underlying neural mechanisms involved in human-AI interactions by extracting pre-stimulus oscillatory activity and examining its modulatory effects on post-stimulus evoked responses. As AI-generated visual content becomes increasingly prevalent in everyday life, these findings contribute to a growing understanding of how beliefs and expectations about such content shape human perception and cognition.

## Acknowledgements

This work was supported by the cooperation project between the Max Planck Society and the Fraunhofer Gesellschaft (grant: project NEUROHUM), as well as by the Deutsche Forschungsgemeinschaft (DFG, German Research Foundation) under Germany’s Excellence Strategy—EXC 2002/1 “Science of Intelligence” (project number 390523135) and grant AB277/6 awarded to Rasha Abdel Rahman. Additional funding was provided by the German Federal Ministry of Education and Research, Technology, and Space (BMFTR) under grants 13GW0206, 13GW0488, and 16SV9156, and by the DFG under grants 502864329 and 542559580 awarded to Michael Gaebler. We also thank other members of project NEUROHUM for their suggestions and all the participants contributing to this study.

## Declaration of competing interest

The authors declare that they have no known competing financial interests or personal relationships that could have appeared to influence the work reported in this paper.

## Data availability

The datasets acquired during the study are available to researchers upon reasonable request to the corresponding author and with appropriate institutional review board approval. The preprocessed data sets from the previous article (Eiserbeck et al., 2023) are available here: https://osf.io/4y8gs/overview.

## Supplementary S1: ERP waveforms

**Figure S1:**
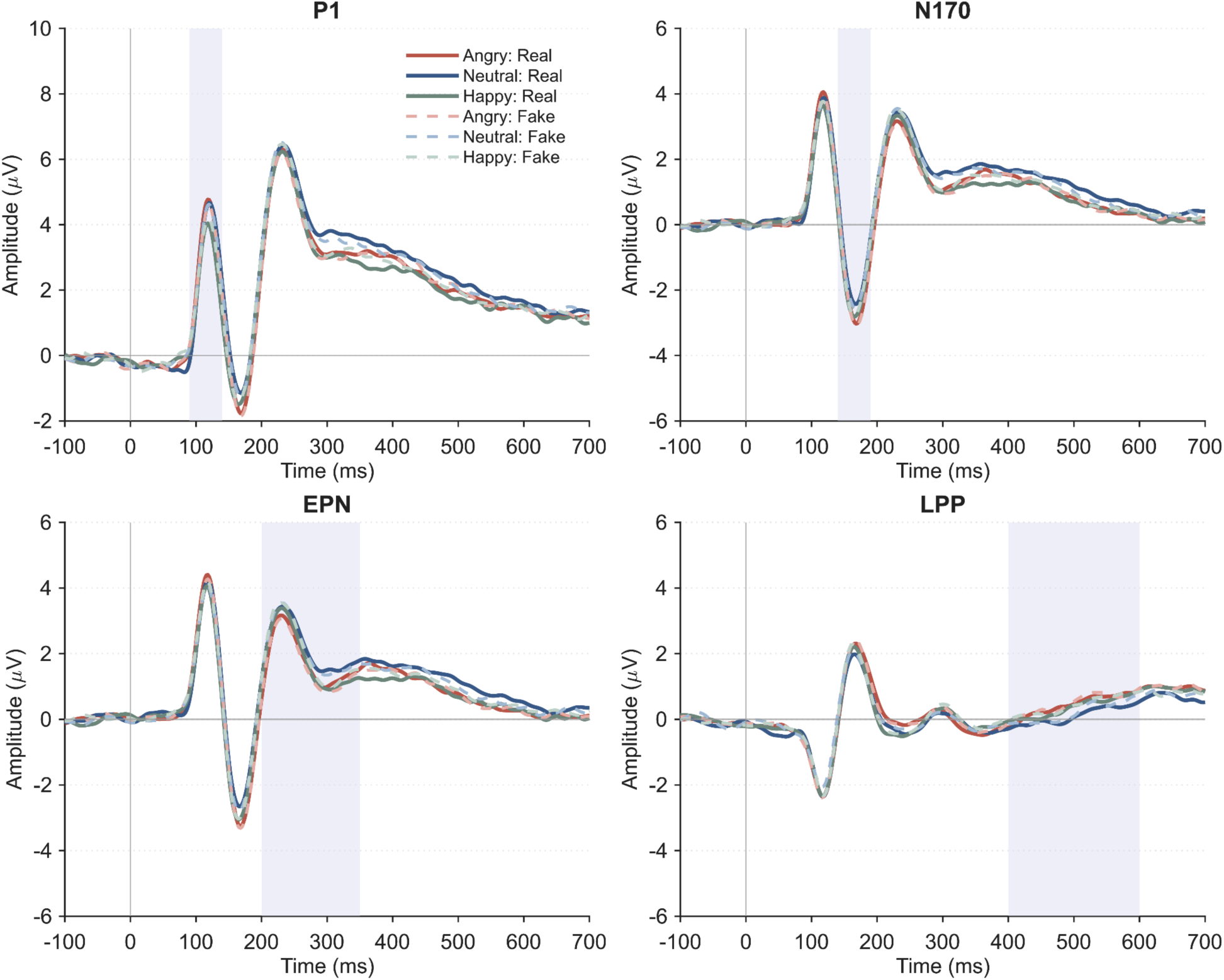
Grand-average ERP waveforms for the four components. Gray shaded areas indicate the specific time windows used for calculating the amplitudes of each component. For details regarding the specific electrode clusters utilized for each ERP, refer to the Methods section.

## Supplementary S2: ERP comparisons

**Table S2:** Summary of Statistical Comparisons of ERP Amplitudes.

|  | “REAL” |  | “FAKE” |  |
| --- | --- | --- | --- | --- |
| | $A_{real,happy}$ | $A_{real,angry}$ | $A_{fake,happy}$ | $A_{fake,angry}$ |
| **P100** | -0.44±0.85** | 0.10±0.65 | -0.28±0.74* | -0.04±0.86 |
| **N170** | -0.38±0.51*** | -0.41±0.64** | -0.21±0.54 | -0.43±0.59*** |
| **EPN** | -0.33±0.41*** | -0.38±0.50*** | -0.13±0.41 | -0.48±0.53*** |
| **LPP** | 0.25±0.55* | 0.33±0.41*** | 0.22±0.36* | 0.28±0.39*** |

$A_{instr,emo}$ were evaluated by computing the difference in ERP amplitudes ( $\mu V$ ) between emotional facial stimuli (Happy & Angry) and neutral stimuli. Values are presented as mean ± standard deviation ( $M \pm SD$ ) for the corresponding $A_{instr,emo}$ . Significance levels of comparison results were acquired after applying paired-samples t-tests between emotional conditions and neutral emotions (\*: $p < 0.05$ , \*\*: $p < 0.01$ , \*\*\*: $p < 0.001$ ).

## Notes

### Competing Interest Statement

The authors have declared no competing interest.

